# Cannabidiol administration reduces the expression of genes involved in mitochondrial electron transport chain and ribosome biogenesis in mice CA1 neurons

**DOI:** 10.1101/2023.07.10.548420

**Authors:** João P. D. Machado, Valéria de Almeida, Antonio W. Zuardi, Jaime E. C. Hallak, José A. Crippa, André S. Vieira

**Affiliations:** Laboratory of Electrophysiology, Neurobiology and Behaviour, Dept. Functional and Structural Biology, Institute of Biology, University of Campinas; Laboratory of Neuroproteomics, Dept. Biochemistry and Tissue Biology, Institute of Biology, University of Campinas; Brazilian Institute of Neuroscience and Neurotechnology (BRAINN), Campinas, São Paulo, Brazil; Department of Neuroscience and Behavior, Ribeirão Preto Medical School, University of São Paulo, Ribeirão Preto, Brazil; National Institute for Science and Technology – Translational Medicine, Brazil

**Author notes:** **Correspondence to:** André Schwambach Vieira, Ph.D., Department of Structural and Functional Biology, Institute of Biology, Rua Monteiro Lobato, 255 - Bloco J - 1 piso - Sl 16, Cidade Universitária “Zeferino Vaz”, University of Campinas – UNICAMP, Campinas - SP, Brazil, 13083-862.

**Keywords:** CBD, RNA-Seq, transcriptomics, laser-capture microdissection, hippocampal subfields

## Abstract

**Background:** Cannabidiol (CBD), one of the main cannabinoids present in the female flowers of *Cannabis sativa*, has been a therapeutic alternative for a plurality of disorders. Previous investigation has already provided insights into the CBD molecular mechanism, however, there is no transcriptome data for CBD effects on hippocampal subfields. Here, we explore the transcriptomic changes in dorsal and ventral CA1 of adult mice hippocampus after 100 mg/kg of CBD administration (i.p.) for one or seven consecutive days.

**Methods:** C57BL/6JUnib mice were divided into 4 groups treated with either vehicle or CBD for 1 or 7 days. The collected brains were sectioned and the hippocampal subregions were laser microdissected for RNA-Seq analysis. Data alignment, quantification and analysis were conducted with the STAR Aligner/DESeq2/clusterProfiler R-package pipeline.

**Results:** We found changes in gene expression in CA1 neurons after single and multiple CBD administrations. Furthermore, the enrichment analysis of differentially expressed genes following 7 days of CBD administration indicates a widespread decrease in the expression levels of electron transport chain and ribosome biogenesis transcripts, while chromatin modifications and synapse organization transcripts were increased.

**Conclusion:** This dataset provides a significant contribution toward advancing our comprehension of the mechanisms responsible for CBD effects on hippocampal neurons. The findings suggest that CBD prompts a significant reduction in energy metabolism genes and the protein translation machinery in CA1 neurons.

**SIGNIFICANT OUTCOMES:** We identified distinct changes in gene expression of CA1 neurons following both single and multiple administrations of CBD. This highlights the molecular impact of CBD on hippocampal neurons and expands our understanding of its mechanisms of action. We revealed that repeated CBD administration led to a greater number of gene expression alterations compared to a single administration, emphasizing the importance of treatment frequency in modulating gene expression. We found that daily CBD administration for seven days resulted in the downregulation of genes related to energy metabolism and protein synthesis/degradation, while genes involved in chromatin regulation and synapse organization were upregulated. These specific gene expression changes shed light on potential cellular effects and molecular mechanisms underlying CBD’s actions in the hippocampus.

**LIMITATIONS:** One limitation of this study is its reliance on animal models, specifically C57BL/6JUnib mice, which may not fully reflect human responses to CBD. Additionally, the study primarily investigated the effects of CBD under healthy conditions and did not directly address its therapeutic effects for specific disorders or conditions. Thus, the clinical relevance and applicability of the findings to therapeutic interventions remain to be determined.

## 1 INTRODUCTION

Cannabidiol (CBD) is a naturally occurring compound found in female flowers of *Cannabis sativa*, an ancient plant with a long history of agriculture by humanity (Ren et al., 2021). In Brazil, there are considerable challenges when it comes to the use and investigation of cannabis. The presence of strict drug policies and a history of drug-related violence creates obstacles, restricting access to cannabis and hinder scientific research. These limitations prevent both researchers and patients from fully exploring the potential of cannabis as a treatment option and comprehending the intricate molecular mechanisms and gene activation associated with cannabis compounds, like CBD.

CBD has a complex pharmacodynamic profile and most pharmacological targets are enzymes, membrane transporters, ionic receptors, and channels (Bih et al., 2015). Regardless of being a phytocannabinoid and having a structural similarity to Δ9-THC, CBD exhibits low affinity to the CB1 receptor orthosteric site (McPartland et al., 2007). This may be a possible explanation for the lack of CBD intoxicating effects. Preclinical research has suggested that CBD may have potential therapeutic effects as an antiepileptic, antioxidant, antipsychotic, and anxiolytic agent (Elsaid et al., 2019). Clinical studies have further established the safety and efficacy of CBD in controlling symptoms of seizures in childhood epilepsies (Devinsky et al., 2018). Despite these promising findings, the underlying molecular mechanisms and gene activation associated with CBD in brain cells are still not fully understood.

One of the brain regions most affected by a plurality of psychiatric and neurological disorders such as depression, schizophrenia, Alzheimer, and epilepsy is the hippocampus (Anand and Dhikav, 2012). The hippocampus is a structure segmented into four sub-regions (CA1, CA2, CA3, DG) that form a complex circuitry involved mainly in learning and memory processes (Fanselow and Dong, 2010; Squire et al., 2015). Each of these subregions has distinct molecular mechanisms and neuronal connections, thus selecting cellular populations precisely helps to specifically explore hippocampal functionality (Machado et al., 2022).

In the present study, we used the Laser Microdissection Technique (LCM - Laser Capture Microdissection) in conjunction with RNA-Seq to select particular cell populations, dorsal and ventral CA1, preserving specific expression characteristics of such regions. With these strategies, we explore the effect of CBD on the transcriptome of dorsal and ventral CA1 after intraperitoneal administration for 1 or 7 days.

## 2 METHODS

### 2.1 Animals and Drugs

Three-month-old males C57BL/6 Unib were housed in a ventilated environment (12 h/12 h light cycle) with *ad libitum* access to standard rodent chow and water. All procedures were executed in compliance with ethical standards for animal experimentation at the University of Campinas-UNICAMP (Brazilian federal law 11.794 (10/08/2008 - Animal Use Ethics Committee protocol 2903-1). We used the RNASeqPower package in the R environment to determine the appropriate sample size (https://bioconductor.org/packages/release/bioc/html/RNASeqPower.html). Mice were divided into four experimental groups (n=5): CR1 - 1 day of NaCl (0.15 M) administration; CBD1 - 1 day of treatment with CBD (100 mg/kg); CR7 - 7 days of NaCl (0.15 M) administration; CBD7 - 7 days of treatment with CBD (100 mg/kg). All solutions were made with NaCl at 98% and Tween 80 at 2% (Sigma-Aldrich, UK) and administered i.p. in a volume of 1 mL/kg.

After 24 hours from the last i.p. administration of CBD, the animals were anesthetized with isoflurane (2% isoflurane, 98% oxygen at 1 liter/min). Once deep anesthesia was confirmed by the absence of corneal reflex, the animals were decapitated using a small animal guillotine, and their brains were immediately collected and frozen at -55°C. The previously collected brains were processed using a cryostat (Leica Biosystems - Wetzlar, Germany) to produce histological slices of 60µm for each animal. The beginning of the slice collection was at the visualization of the third ventricle, and the end of the collection was after the disappearance of the granular cell layer of the DG. The slices were collected on PEN membrane-coated glass slides (Life Technologies®, Thermo Fisher Scientific - Waltham, USA), immediately stained with Cresyl Violet, dehydrated through an increasing alcohol series (70% ethanol at -20ºC for 2 minutes; 1% Cresyl Violet acetate for 2 minutes; washed with 70% ethanol and dried at room temperature), and stored at -80°C.

### 2.2 Laser microdissection

For laser microdissection, the previously stored slides were brought to room temperature, placed on the microscope, and delimited with the PALM system (Zeiss® - Jena, Germany). The boundaries of the ventral and dorsal CA1 region were located based on previous studies of the regions (Fanselow and Dong, 2010), and markers found in the *in situ hybridization* images of the Allen Brain Atlas (http://portal.brain-map.org/). We also followed the hippocampal subfield microdissection methods as described by (Vieira et al., 2016). Subsequently, the previously microdissected ventral and dorsal CA1 subregion were mechanically collected and isolated in plastic microtubes using ophthalmic forceps and a surgical microscope (Zeiss® - Jena, Germany).

### 2.2 Library preparation and Next-generation sequencing

RNA was extracted from the samples using TRIzol (Thermo Fisher Scientific, USA) according to the manufacturer instructions. The quality of the extracted RNA was assessed by the 2100 Bioanalyzer Instrument using Agilent RNA 6000 Pico kit (Agilent, CA, USA) and an average integrity quality number (RIN) of 7 was obtained for all samples. The cDNA libraries were synthesized using TruSeq Stranded Total RNA LT (Illumina®, CA, USA) from 200 ng of RNA, following the manufacturer instructions. Barcoded libraries were pooled and sequenced on a HiSeq® 2500 (Illumina®, CA, USA) in High Output mode, generating 100-bp paired-end sequences. On average, 20 million paired-end reads were obtained per sample, totaling 794.682.591 reads across all samples. The gene expression and statistical analysis were performed in DESEQ2 using read counts per gene and average sequence alignment of 69,89%.

### 2.3 Data processing and differentially expressed genes (DEGs)

We aligned all sequenced reads with the *Mus musculus* genome (GRCm38.96) using the StarAligner 2.6 program (https://github.com/alexdobin/STAR, RRID: SCR_004463) (Dobin et al., 2013). To normalize the data and perform statistical analysis of differentially expressed genes, the DESeq2 package (http://bioconductor.org/packages/release/bioc/html/DESeq2.html) was used, which is implemented in the R platform (https://cran.r-project.org/bin/) (Love et al., 2014). Statistical analysis and enrichment analysis were carried out using DESeq2 and clusterProfiler package (https://bioconductor.org/packages/release/bioc/html/clusterProfiler.html, RRID: SCR_016884) (Wu et al., 2021). Enrichment analysis classified the functional profiles into KEGG Pathways and GO Terms - Biological Processes (BP). Differential gene expression was considered significant when adjusted p-value < 0.05. The significance of terms and pathways in the enrichment analysis was determined using a p-value < 0.05 (after adjustment for multiple comparisons using the Bonferroni Test).

## 3 RESULTS

### 3.1 Identification of DEGs and Samples Visualization

A principal component analysis (PCA) was conducted for dorsal CA1 (dCBD1vsCR1 and dCBD7vsCR7) and ventral CA1 (vCBD1vsCR1 and vCBD7vsCR7) (Figures 1A-D). Both dorsal and ventral CA1 samples revealed an overlap of the CBD1 and CR1, while the CBD7 and CR7 formed distinct groups. We obtained the following number of DEGs (adjusted p < 0,05): 6 (dCBD1vsCR1), 1559 (dCBD7vsCR7), 299 (vCBD1vsCR1), 2924 (vCBD7vsCR7). The number of upregulated and downregulated genes is represented in Figure 1E. For a complete list of DEGs refer to Supplementary Table 1. Finally, the Venn diagram illustrates the common and distinct DEGs among the groups (Figure 1F).

**Figure 1.**
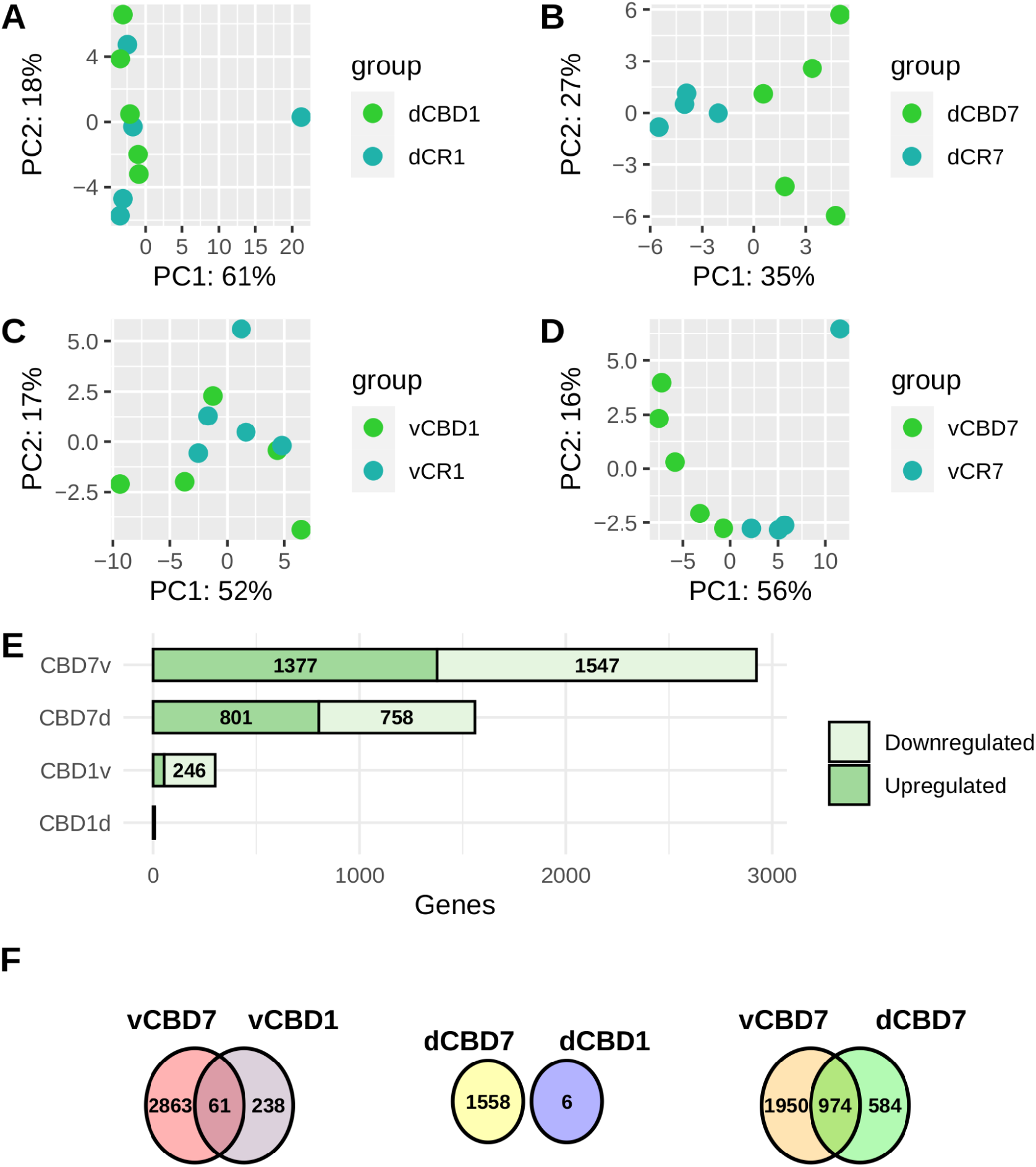
Sample variability and distribution of differentially expressed genes. **(A)** Principal component analysis of gene expression data from dorsal CBD1. **(B)** Principal component analysis of gene expression data from dorsal CBD7. **(C)** Principal component analysis of gene expression data from ventral CBD1. **(D)** Principal component analysis of gene expression data from ventral CBD1. **(E)** Barplot displaying the number of differentially expressed transcripts identified in each ventral and dorsal CBD analysis. **(F)** Venn diagram representing common and unique genes expressed between CBD groups.

### 3.2 GO and pathways

#### 3.2.1 CBD effect on dorsal CA1

GO and KEGG analysis identified a total of 87 significantly (p.adjust < 0.05) enriched biological processes (BPs) and 26 significantly (p.adjust < 0.05) enriched pathways for dCBD7 (Supplementary Table 2). We also performed GO enrichment using upregulated and downregulated genes to determine the specific transcriptional change of biological functions (Figure 2A - 3A). For the dCBD1 group, there was only one significant biological process (p.adjust < 0.05) related to “*skeletal muscle cell differentiation*”.

**Figure 2.**
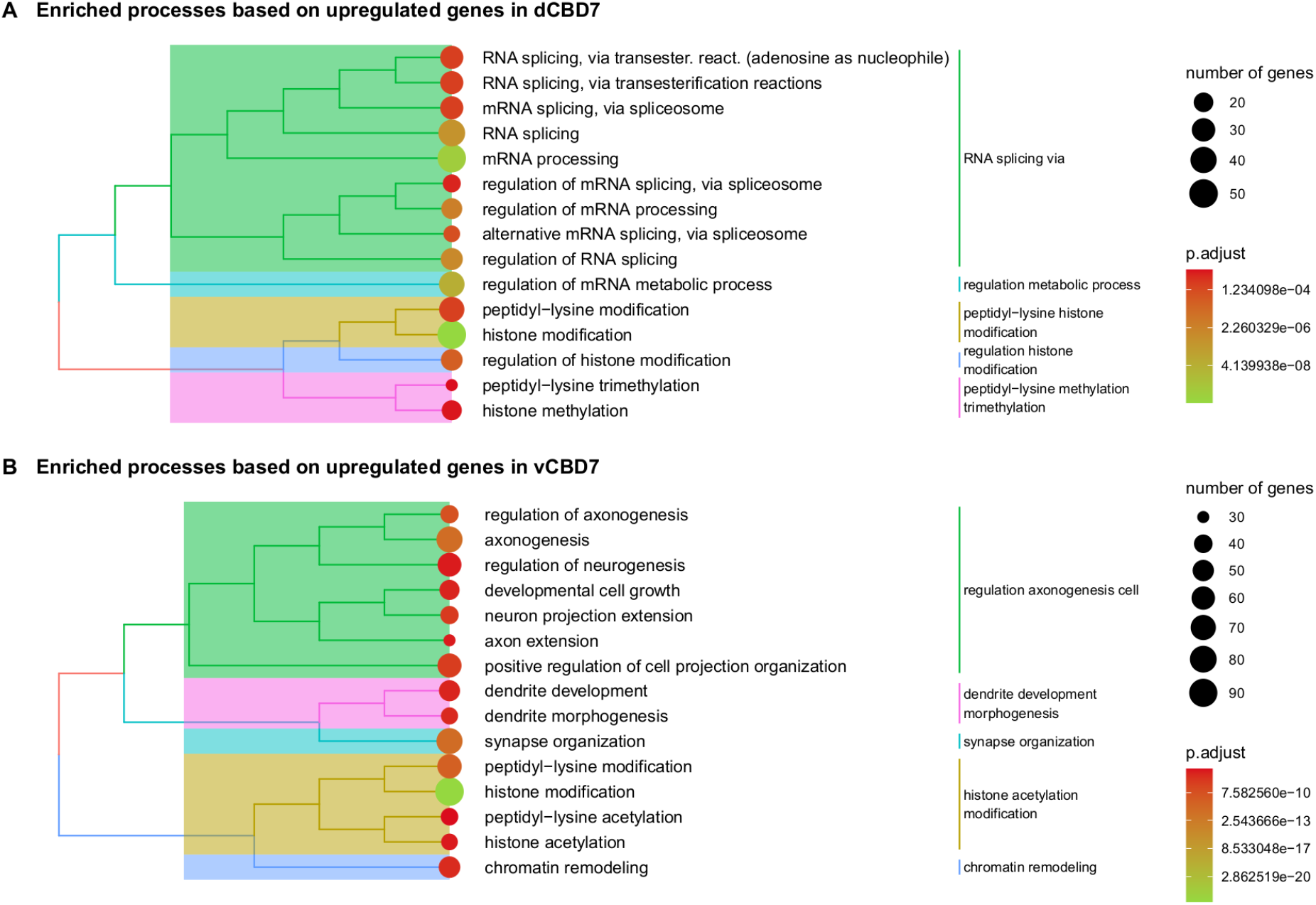
Enriched biological processes for differentially expressed genes in CBD7. **(A)** Top enriched biological processes for upregulated genes in dorsal CBD7. **(B**) Top enriched biological processes for upregulated genes in ventral CBD7. All displayed biological processes reached an adj. p-value < .05.

**Figure 3.**
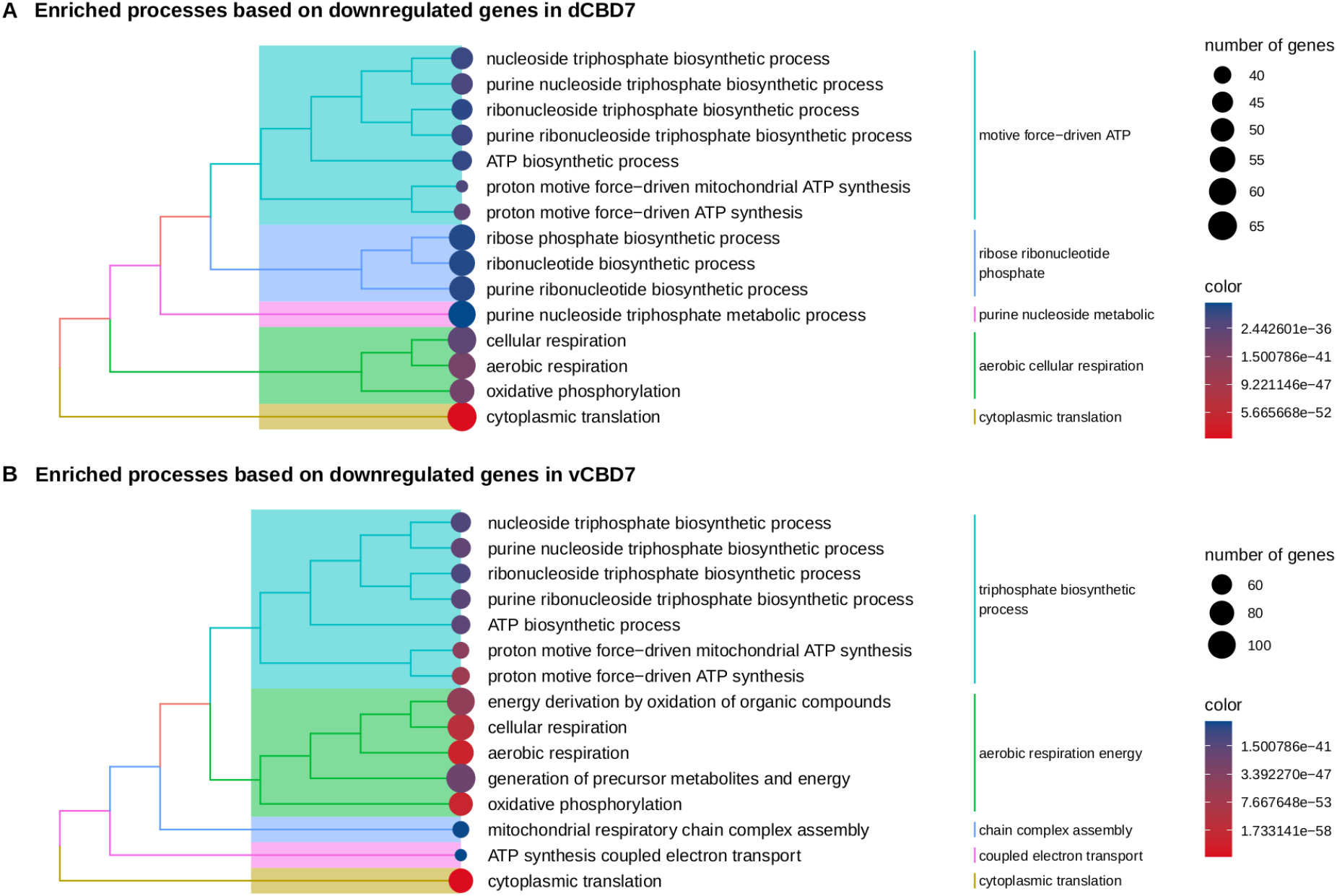
Enriched biological processes for differentially expressed genes in CBD7. **(A)** Top enriched biological processes for downregulated genes in dorsal CBD7. **(B)** Top enriched biological processes for downregulated genes in ventral CBD7. All displayed biological processes reached an adj. p-value < .05.

#### 3.2.2 CBD effect on ventral CA1

GO and KEGG analysis identified a total of 87 significantly (p.adjust < 0.05) enriched biological processes (BPs) and 26 significantly (p.adjust < 0.05) enriched pathways for vCBD7 (Supplementary Table 2). We also performed GO enrichment using upregulated and downregulated genes to determine the specific transcriptional change of biological functions (Figure 2B - 3B). For the ventral CBD1 group, there was only one significant biological process (p.adjust < 0.05) related to “*proteasome-mediated ubiquitin-dependent protein catabolic process*”.

#### 3.2.3 CBD effect on the ribosome and oxidative phosphorylation (OXPHOS) gene expression

The majority of genes enriched in biological processes exhibiting reduced expression are involved in energy metabolism, specifically in “mitochondrion organization,” “ATP metabolic process,” “oxidative phosphorylation,” and “electron transport chain” (Figure 5 A-B). The remaining genes are implicated in protein translation, including “ribonucleoprotein complex biogenesis,” “cytoplasmic translation,” and “ribosome assembly” (Figure 5 C-D). Additionally, the enriched pathways confirmed “Ribosome” and “Oxidative phosphorylation” as the most significant pathways, while also highlighting “Proteasome,” “RNA polymerase,” “Protein export,” and “Citrate cycle (TCA cycle)” as enriched pathways (Supplementary Table 2).

## 4 DISCUSSION

In this study, we investigated alterations in gene expression using LCM/RNA-Seq among distinct hippocampal neuron subtypes following CBD administration. We aimed to unveil the impact of CBD on transcriptome modifications and pertinent biological processes and pathways associated with these changes. Additionally, we aimed to differentiate between effects within different hippocampal regions, dorsal and ventral, across two administration protocols (1-day and 7-day intervals). Our findings revealed, for the first time, a reduction in the expression of several genes linked to the respiratory chain and ribosomal subunits in an animal model, accompanied by an increase in the expression of genes related to histone regulation. Furthermore, neural projection formation genes were specifically upregulated in the ventral CA1, while RNA splicing genes were specifically upregulated in the dorsal CA1. Overall, these results contribute significantly to advancing our knowledge of the effects of CBD on gene expression in hippocampal neurons, thus elucidating the molecular mechanisms underlying its effects.

### 4.1 Expression alteration comparing administration frequency

Our study reveals a significant difference in the differentially expressed genes observed between the two durations of CBD treatments. Prior research has noted behavioral and molecular distinctions between single and chronic CBD administrations (50 mg/kg) in C57BL/6JArc mice (Darweesh et al., 2020; Long et al., 2010). Long et al. (2010) observed that the anxiolytic and antipsychotic effects of CBD are exclusively achieved after chronic administration in the model used by the authors. Chronic administration of CBD alters other pharmacokinetic pathways of carbamazepine metabolism beyond CYP3A, which is exclusively altered in a single dose (Darweesh et al., 2020). CBD ability to bind to multiple molecular targets may play a role in producing different physiological effects. Furthermore, binding duration is critical in determining the effectiveness and potency of pharmacological responses of several targets affected by CBD affinity and bioavailability.

While some similarities in molecular effects of dosages are observed, there are discrepancies in administration frequency (Long et al., 2010; Valvassori et al., 2013). CBD doses of 60 mg/kg resulted in increased mitochondrial activity of complexes I, II, III, and IV in both acute and chronic administration in rats hippocampus, prefrontal cortex, and cerebral cortex (Valvassori et al., 2013). On the other hand, reducing the dosage to 15mg/kg led to no response in a single dose of CBD, but increased activity of complexes II and IV in the hippocampus and striatum of the chronic dosage group (Valvassori et al., 2013). Although previous studies have demonstrated similarities and differences, we are the first to report a significant number of differentially expressed genes in two administration frequencies. Therefore, our data clarifies that the transcriptome of dorsal and ventral CA1 neurons undergoes more significant alteration after 7 days of CBD administration than a single dose.

### 4.2 Reduction of expression of genes from the respiratory chain and tricarboxylic acid cycle

We observed a reduction in transcriptional levels of all mitochondrial respiratory chain complexes both in ventral and dorsal CA1 after administering 100 mg/kg CBD for 7 days **(Figure 4 A-B)**. The CBD ability to decrease oxidative metabolism in tissue and isolated mitochondria has long been known (Chiu et al., 1975). *In vitro* studies have shown that CBD can reduce oxygen consumption and mitochondrial complex activity (Fišar et al., 2014; Singh et al., 2015). However, Valvassori et al. (2013) observed an increased mitochondrial complex activity in the rats’ hippocampus after administering 60 mg/kg i.p of CBD for 14 consecutive days. In this context, the methodology with the most similarity is Valvassori study, although our data clearly show a reduction in the expression of mitochondrial complexes after CBD administration. It should be noted that studies on the specific effects of CBD on mitochondria differ in model, dosage, and administration method. In addition, the reduced expression of respiratory chain complexes highlighted in our data may not necessarily indicate compromised mitochondrial function due to limiting factors of respiration rate and compensatory mechanisms (Brand and Nicholls, 2011).

**Figure 4.**
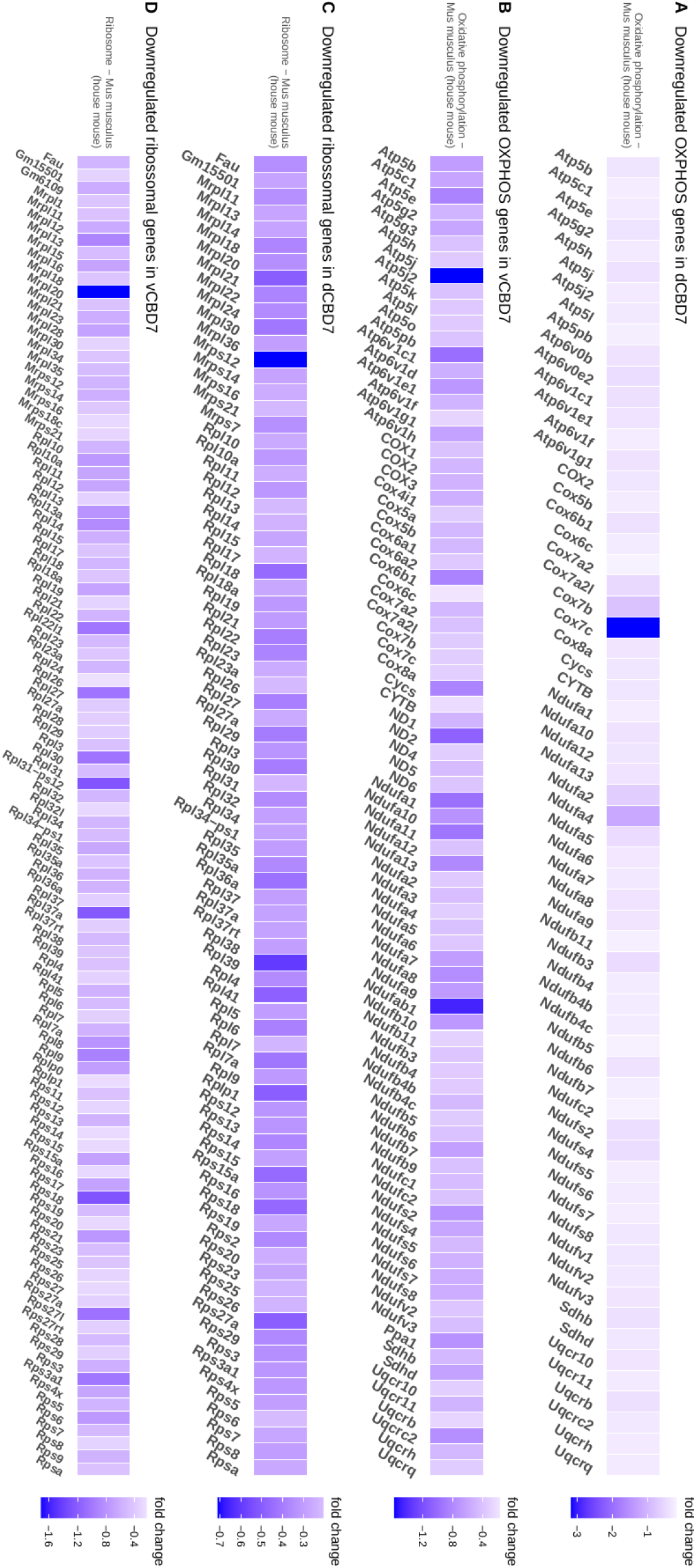
Differentially expressed genes in CBD7 for enriched ribosome and OXPHOS processes. **(A)** Downregulated genes in dorsal CA1 related to OXPHOS. **(B)** Downregulated genes in ventral CA1 related to OXPHOS. **(C)** Downregulated genes in dorsal CA1 related to ribosomes. **(D)** Downregulated genes in ventral CA1 related to ribosomes. All displayed genes reached an adj. p-value < .05.

The previously mentioned studies’ findings are crucial, as they indicate a biphasic effect of CBD on mitochondrial respiration. Evidence suggests that at lower or moderate dosages, CBD enhances mitochondrial respiration, whereas, at higher concentrations, it inhibits them. This observation underscores the significance of the dose-dependent effects of CBD on mitochondrial function and emphasizes the need for further functional studies underlying the mechanisms. Other endogenous and exogenous CB1 ligands induced a biphasic *in vitro* change in some mitochondrial complexes (Athanasiou et al., 2007). Nevertheless, a direct comparison of respiratory chain gene activity and expression levels after different CBD dosages in animal models is necessary to fully understand the effects on mitochondrial function.

We are the first to observe that CBD administration reduced genes from the subunits of enzymes in the tricarboxylic acid cycle (TCA) in ventral and dorsal CA1, such as *Idh3a* (*isocitrate dehydrogenase 3 (NAD+) alpha*), *Suclg1* (*succinate-CoA ligase, GDP-forming, alpha subunit*) and *Mdh1* (*malate dehydrogenase 1*). A study with synthetic cannabinoids had already reported an increase in α-ketoglutarate levels, indicating inhibition of the α-ketoglutarate dehydrogenase enzyme (Dando et al., 2013). However, in the case of CBD, we only observed a reduction in the genes of isocitrate, succinyl-CoA, succinate, and malate dehydrogenase enzymes. In addition, another study noted a decrease in the abundance of most of the TCA cycle metabolites in CBD-treated cells (Guard et al., 2022). Therefore, due to the marked alteration in the gene expression of the TCA cycle enzyme, it becomes more likely that CBD alters mitochondrial activity.

Despite having a low affinity for CB1 receptors, CBD is known to inhibit enzymes responsible for endocannabinoid metabolism, leading to increased activation of these receptors. CB1 receptor activation by endocannabinoids can reduce the expression of genes involved in mitochondrial biogenesis and oxygen consumption in white adipocytes of mice (Tedesco et al., 2010). The direct inhibition of complex I activity in CA1 by the endocannabinoid system contributes to inhibiting inhibitory postsynaptic potentials through a process known as depolarization-induced suppression of inhibition (Bénard et al., 2012). In addition to the decrease in the expression of genes associated with the respiratory chain and TCA cycle, our analysis revealed the enrichment of several biological processes related to axon and dendrite formation following a 7-day administration of CBD in ventral CA1. The endocannabinoid system inhibition of the respiratory chain and the resulting decrease in ATP production may represent a possible mechanism for efficiently modulating high-energy-demand synaptic processes such as neurotransmitter release and neuroplasticity (Bénard et al., 2012). Thus, the observed decrease in the expression of crucial mitochondrial genes and increase in genes regulating synaptic modulation may be associated with endocannabinoid signaling. Nevertheless, further investigations are necessary to elucidate the underlying molecular mechanisms and confirm this hypothesis.

### 4.3 Reduction of gene expression of ribosomal and proteasomes subunits

Ribosomes are essential for protein synthesis, and a decrease in their abundance has been demonstrated to reduce protein synthesis (Jakubovic and McGeer, 1972). Moreover, the availability of ATP is critical for ribosomal RNA transcription and can lead to ribosome formation and also translation (Murray et al., 2003). Cell regulates the rate of protein synthesis by maintaining a balance between ATP consumption and ribosomal transcription. Early studies have shown that Δ9-THC application causes a reduction of ribosome number (Hattori et al., 1972). We are the first to report a significant reduction in gene expression of several ribosomal subunit constituents in both ventral and dorsal CA1 regions after 7 days of CBD administration **(Figure 4 C-D)**. In this way, the reduction in ribosomal subunit expression identified by our study may correspond to the state of mitochondrial activity and protein synthesis but further investigation is required.

Maintaining the balance between protein synthesis and degradation is crucial for cellular homeostasis (Xolalpa et al., 2013). One of the mechanisms capable of modifying and recycling proteins in cells is the proteasomes. Our results also show several proteasome subunits downregulated after 7 days of CBD administration. Furthermore, the differentially expressed genes found after a single application of CBD may offer an initial explanation for the observed effect on the proteasome following repeated administration, given the relationship between ubiquitin and the proteasome in protein degradation. Reduction in genes involved in the ubiquitin/proteasome system may reflect a lower protein synthesis, as fewer proteins are degraded or modified. However, further investigation is required to validate the underlying cellular mechanisms involved in protein synthesis and degradation following CBD administration.

### 4.4 Increase in expression of genes regulating small GTPases and neuroplasticity

Small GTPases are a group of enzymes involved in various cellular processes, and our findings show statistically significant terms related to them. Cannabinoids are known to alter the signaling of some small GTPases through CB1 and CB2 receptors (Duman et al., 2015; Kurihara et al., 2006), and observing the effects of repeated CBD administration on these molecules provides insight into the intracellular responses and possible repercussions of this signaling. We are the first to find altered expression of genes that encode regulatory proteins of GTPases after administration of 100 mg/kg of CBD for 7 days, such as *Arhgap17* (*Rho GTPase-activating protein 17*), *Arhgap 21* (*Rho GTPase-activating protein 21*), *Arhgap 39* (*Rho GTPase-activating protein 39*), *Arhgef1* (*Rho guanine nucleotide exchange factor 1*), *Arhgef2* (*Rho guanine nucleotide exchange factor 2*), *Myo9b* (*Myosin ixb*), *Akap13* (*A-kinase anchor protein 13*). The small GTPases of the Rho family are known for their important regulatory effects on the organization of the postsynaptic density and actin cytoskeleton (Stankiewicz and Linseman, 2014). Thus, as the activity of small GTPases depends on their modulators, the presence of CBD may modify these signaling pathways and consequently alter the cytoskeleton and synaptic plasticity.

Neural plasticity is the ability of neural networks to transform through the growth and reorganization of synapses. Our findings reveal that CBD increases the expression of genes involved in synapse organization and dendrite and axon formation only in the ventral sub-region of CA1 in the hippocampus. Previous studies have shown that some phytocannabinoids can modify the number of microtubules present in PC12 cells *in vitro (Tahir et al., 1992*). We observed that 100 mg/kg CBD for seven days increases the expression of genes such as *Actn1* (*Alpha-actinin-1*), *Flna* (*Filamin-A*), *Synpo* (*Synaptopodin*), *Shank1/Shank2* (*SH3 and multiple ankyrin repeat domains protein*), and *Itsn1* (*Intersectin-1*). These proteins are responsible for mediating the morphology of dendritic spines and the organization of the postsynaptic density (Muñoz-Lasso et al., 2020). Changes in genes that interact with the actin cytoskeleton can influence synapse formation due to greater stabilization in neuronal process development (Muñoz-Lasso et al., 2020; Sarowar and Grabrucker, 2016). Therefore, the effect of CBD on the neuronal cytoskeleton in the hippocampus may induce changes in dendritic morphology, resulting in the development or shrinkage of dendritic spines and promoting plasticity between circuits.

One important aspect of neuroplasticity and nervous system formation is the guided encounter through chemical signaling between the axonal projections of the presynaptic neuron and the dendritic tree of the postsynaptic neuron. Semaphorins, plexins, and neuropilins are among the numerous signaling molecules and receptors that work together to regulate axonal guidance (Christie et al., 2021). The data presented in this study show an increase in the expression of these molecules, such as *Sema4d* (*Semaphorin-4D*), *Sema4f* (*Semaphorin-4F*), *Sema5a* (*Semaphorin-5A*), *Plxna2* (*Plexin-A2*), *Plxna3* (*Plexin-A3*), *Plxnb2* (*Plexin-B2*), and *Nrp2* (*Neuropilin-2*), in the ventral CA1 region after the administration of 100 mg/kg of CBD for 7 days. The interaction between these molecules, such as *Sema5d*/*Plexna2*, inhibits the formation of excitatory synapses in mouse hippocampal cells (Duan et al., 2014). Furthermore, the addition of *Sema4d* in the CA1 subregion of mice promotes the formation of inhibitory synapses and suppresses seizures in models that generate crises via electrical and chemical stimuli (Acker et al., 2018). CBD has long been known to reduce the number of seizures and increase the seizure threshold, especially in animal models (Consroe et al., 1982). Therefore, these results may provide clues on how CBD regulates molecules that are responsible for the plasticity of inhibitory and excitatory synapses in ventral CA1, as well as to better understand possible mechanisms that control seizures.

Changes in gene expression that control axon and dendrite growth are a clue to potential alterations in long-term depression/potentiation (LTD/LTP). In the hippocampus, the endocannabinoid system has a profound effect on LTD/LTP and modulates synaptic structure and activity plasticity (Robledo-Menendez et al., 2022). Treatment with Δ9-THC for seven days completely abolishes the LTP of CA1 pyramidal neurons (Hoffman and Lupica, 2013). Moreover, the Δ9-THC-induced decrease in LTP is prevented by pharmacological inhibition or exclusion of the CB1 receptor (Hoffman and Lupica, 2013). On the other hand, the ability of CBD to increase the LTP generation in the CA1 region has also been described *in vitro* mouse hippocampus (Maggio et al., 2018). An increased presence of F-actin is a possible marker for LTP (Fukazawa et al., 2003). Although we did not find it differentially expressed, we observed some genes in our ventral CA1 data that were identified through high-performance sequencing of a late phase of LTP (Chen et al., 2017), such as *Cntnap2* (*contactin associated protein-like 2*), *Vcl* (*vinculin*), *Pcdhga9* (*protocadherin gamma subfamily A, 9*), *Col11a1* (*collagen, type XI, alpha 1*), *Col4a1* (*collagen, type IV, alpha 1*), *Flnb* (*filamin, beta*), *Micall1* (*microtubule associated monooxygenase, calponin and LIM domain containing -like 1*) e *Numa1* (*nuclear mitotic apparatus protein 1*). Although it is uncertain whether LTP/LTD events were present in our study, our findings indicate that repeated CBD administration can increase the expression of genes involved in neuroplasticity processes, such as axon and dendrite development. This sheds light on the involvement of the endocannabinoid system in neuroplasticity and suggests the potential of CBD as a therapeutic intervention for related dysfunctions.

Defects in processes involving neuroplasticity are well-known in the pathophysiology of mood disorders and diseases (Dorszewska et al., 2020; Post, 1992). A single administration of 30 mg/kg of CBD, despite alleviating depressive-like behaviors, was unable to restore decreased dendritic spine density in the ventral CA1 region after the stress-induced depression model (Ma et al., 2021). However, repeated CBD treatment promotes dendritic spine remodeling in the hippocampus of chronically stressed mice (Fogaça et al., 2018). This CBD effect was abolished with the use of CB1/CB2 antagonists, supporting the possibility of synaptic density alteration being mediated by endocannabinoid receptors (Fogaça et al., 2018). Similar to our findings, changes in the expression of plasticity and cytoskeleton genes were only observed with repeated doses, not with a single dose. Moreover, a single dose did not cause the same magnitude of gene expression modification. Thus, the behavioral effects obtained with a single CBD dose must be related to other processes beyond gene expression, such as the excitatory/inhibitory balance of specific circuits in the brain, and changes in plasticity can only be achieved with repeated CBD administration.

### 4.5 Increase in the expression of chromatin regulatory genes

Epigenetic modifications are alterations in chromatin organization induced by external factors, which alter gene expression without affecting the underlying DNA sequence (Rose & Klose, 2014). CBD can increase methylation levels in human keratinocytes *in vitro* by upregulating the expression of *Dnmt1* (Pucci et al., 2013). Furthermore, CBD (10 mg/kg) administration induced antidepressant-like effects and reduced global methylation levels in the prefrontal cortex and hippocampus of mice exposed to the forced swim test (Sales et al., 2020). CBD also modifies DNMT enzyme activity in the prefrontal cortex, and these changes are associated with altered DNA methylation levels and antidepressant-like effects (Sales et al., 2020). Previous studies have shown the importance of DNA methylation, particularly through DNMTs, for neuronal plasticity processes like learning and memory (Feng et al., 2010; Levenson et al., 2006). After 7 days of CBD administration, we observed an upregulation of *Dnmt3a (DNA methyltransferase 3A)* in both dorsal and ventral CA1 regions. It is noteworthy, as *Dnmt3a* is a DNA methyltransferase known to play a role in regulating dendritic spine growth and behavioral traits (LaPlant et al., 2010). These findings suggest that CBD may impact synapse organization processes through the upregulation of DNMT enzymes.

We are the first to provide evidence that CBD appears to regulate the expression of a wide range of executor enzymes involved in the balance of histone modifications essential for gene expression. After 7 days of CBD administration, we found several upregulated methyltransferases (*Kmt2a, Kmt2b, Kmt2c, Kmt2e, and Nsd3*) responsible for the methylation of lysines present in the histone tail. We also identified several enzymes catalyzing the removal of the methyl group from lysines, known as demethylases (*Kdm2a, Kdm4b, Kdm7a, Kdm6b*). These methylation and demethylation reactions on histones can be depicted as a dynamic process of “writer-reader-eraser” enzymes (Hyun et al., 2017). The “writers’’ execute the modifications, while the “erasers’’ remove them, and the “readers’’ are responsible for finding these altered regions. Here, our findings suggest that the 7-day administration of CBD can modulate gene expression in dorsal and ventral CA1 by upregulating transcripts associated with chromatin and histone modifications, providing new players into the epigenetic mechanisms underlying the effects of CBD.

We also observed an upregulation of genes related to RNA splicing and processing in the dorsal CA1 subregion. Among the upregulated genes, we find spliceosome subunits (*Snrnp70* and *Snrnp48*) and splicing factors or splicing regulatory genes (*Srsf1, Srsf11, Srsf2, Srsf9, Sf3b6, Sf3b5, Khdc4, Puf60* and *RBM22)*. These findings suggest that CBD may have a significant impact on alternative splicing by affecting the composition and function of the spliceosome complex. Moreover, DNA methylation and chromatin modifications have been shown to have a significant impact on alternative splicing and RNA processing (Zhang et al., 2020). Additionally, RNA splicing is a crucial mechanism that increases transcriptional complexity and regulates gene expression (Georgakopoulos-Soares et al., 2022). Alternative RNAs can interact with the translational machinery and affect the recognition of the translation start and also decrease translational efficiency (Georgakopoulos-Soares et al., 2022). Further studies are needed to elucidate the molecular mechanisms underlying these effects in neurons and the potential therapeutic applications of CBD in neurological disorders.

In conclusion, the transcriptome data generated in this study reveals the action of CBD in modifying neuronal expression in dorsal and ventral CA1. The effects observed at different frequencies of administration demonstrate that repeated CBD administration is capable of altering the expression of more genes compared to a single administration. Specifically, a single CBD administration caused a small modification in genes related to ubiquitination, while daily CBD administration for seven days reduced the expression of genes involved in energy metabolism and protein synthesis/degradation, but increased the expression of genes involved in chromatin regulation and synapse organization. These results provide evidence for the cellular effects and molecular mechanisms of CBD under healthy conditions, as well as to understand the potential therapeutic effects of its use. However, it is important to consider that the reduced expression of respiratory chain complexes in our data does not necessarily imply compromised mitochondrial function, as other factors and compensatory mechanisms may be involved. Additionally, new potential signaling pathways and genes that are altered after CBD administration in the dorsal and ventral CA1 region of the mouse hippocampus were identified with high statistical confidence. These findings contribute to a better understanding of how CBD alters gene expression and its influence on possible molecular mechanisms in the hippocampus.

## ACKNOWLEDGEMENTS

We acknowledge the use of artificial intelligence (AI) tools, specifically OpenAI ChatGPT, to refine the quality of the English language used in the present paper. The original English text from the authors was corrected by the AI tool and was still revised by the authors to ensure accuracy and clarity.

## AUTHOR CONTRIBUTIONS

**João P. D. Machado:** Conceptualization, Methodology, Investigation, Visualization, Formal Analysis, Writing—Original Draft. **Valéria de Almeida:** Conceptualization, Methodology, Formal analysis, Writing—Original Draft. **Antonio W. Zuardi:** Conceptualization, Methodology, Formal analysis. **Jaime E. C. Hallak:** Conceptualization, Methodology, Formal analysis. **José A. Crippa:** Conceptualization, Methodology, Formal analysis. **André. S. Vieira**: Conceptualization, Methodology, Project administration, Supervision, Writing—Original Draft.

## Supplementary materials

**Supplementary Table 1** – List of genes differentially expressed in all comparisons.

**Supplementary Table 2** – List of GO and KEGG pathways significantly enriched based on all genes in all comparisons.

